# *Wolbachia* strain *w*Au differs in cellular perturbation and virus inhibition profiles from previously characterised *Wolbachia* strains

**DOI:** 10.1101/2021.09.02.458666

**Authors:** Stephanie M Rainey, Vincent Geoghegan, Daniella A Lefteri, Thomas H Ant, Julien Martinez, Cameron McNamara, Steven P Sinkins

**Author notes:** Authors contributed equally (Stephanie M Rainey, Vincent Geoghegan). Corresponding author Steven P Sinkins. **Email:**. **Author Contributions:** SMR, VG & SPS conceived and planned the experiments. JM created the cell lines. THA created the mosquito lines and carried out dissections for proteomics with SMR. SMR, DAL and CM carried out experiments. VG performed proteomics on midguts, SMR & VG analysed proteomic data. SMR & SPS wrote the manuscript with contributions from all authors. **Competing Interest Statement:** The authors declare no competing interests.

## Abstract

Some strains of the inherited bacterium *Wolbachia* have been shown to be effective at reducing the transmission of dengue and other positive-sense RNA viruses by *Aedes aegypti* in both laboratory and field settings and are being deployed for dengue control. The degree of virus inhibition varies between *Wolbachia* strains; density and tissue tropism can contribute to these differences but there are also indications that this is not the only factor involved: for example, strains *w*Au and *w*AlbA are maintained at similar densities but only *w*Au produces strong dengue inhibition. We previously reported perturbations in lipid transport dynamics, including sequestration of cholesterol in lipid droplets, with strains *w*Mel / *w*MelPop in *Ae*. *aegypti*. Here we show that strain *w*Au does not produce the same cholesterol sequestration phenotype despite displaying strong virus inhibition and moreover, in contrast to *w*Mel, *w*Au antiviral activity was not rescued by cyclodextrin treatment. To further investigate the cellular basis underlying these differences, proteomic analysis of midguts was carried out on *Ae*. *aegypti* lines and revealed that *w*Au-carrying midguts showed a distinct proteome when compared to *Wolbachia*-free, *w*Mel- or *w*AlbA-carrying midguts, in particular with respect to lipid transport and metabolism. The data suggest a possible role for perturbed RNA processing pathways in *w*Au virus inhibition. Together these results indicate that *w*Au shows unique features in its inhibition of arboviruses compared to previously characterized *Wolbachia* strains.

**Author Summary:** *Wolbachia* endosymbionts can block transmission of dengue virus by *Aedes aegypti* mosquitoes, and *Wolbachia* release programs for dengue control are now being undertaken in several countries. Understanding the mechanisms of *Wolbachia*-mediated antiviral activity is important for maximizing the efficacy of this control approach. Using functional and proteomic analyses, this study indicates that different strains of *Wolbachia* perturb cellular functions in diverse ways and display different antiviral profiles. These differences raise the possibility that *Wolbachia* strain switching could be used to counteract viral escape mutations, should they arise and threaten the efficacy of dengue control programmes.

## Introduction

The maternally inherited intracellular symbiotic bacteria *Wolbachia* are common in insects and can spread through insect populations by inducing cytoplasmic incompatibility (CI), a sperm modification that results in a pattern of crossing sterility that gives *Wolbachia*-carrying females a relative fitness advantage (1–3). They are not naturally carried by the mosquito *Aedes aegypti*, the primary vector of the flaviruses dengue (DENV), Zika (ZIKV), chikungunya (CHIKV) and yellow fever (YFV), and the alphavirus chikungunya (CHIKV), which together impose a huge public health burden across the tropics (4,5). However, following lab transfers of various *Wolbachia* strains into this mosquito, some strains can block the transmission of DENV, ZIKV, YFV and CHIKV; *Wolbachia* can also inhibit insect-specific flaviruses, West Nile Virus and experimental model arboviruses such as Semliki Forest Virus (SFV) (6–11).

A number of studies have shown that the intracellular density of *Wolbachia* is an important factor in determining the relative ability of *Wolbachia* strains to inhibit viruses (12–15). However, there have been recent indications that density is not the only factor involved: transfer of the *w*Au and *w*AlbA strains, originating in *Drosophila simulans* and *Aedes albopictus* respectively, into *Ae*. *aegypti* resulted in high intracellular densities in both cases, but *w*AlbA produced only limited antiviral activity against DENV/Semliki Forest Virus and a relatively weak capacity to inhibit ZIKV *in vivo* (8,16). In contrast *w*Au produced extremely efficient virus transmission blocking, with no evidence of any DENV dissemination beyond the midgut (8). Mechanistically, a role has been demonstrated for lipid transport and metabolism in the ability of the *w*Mel / *w*MelPop strains from *Drosophila melanogaster* to inhibit dengue *in vivo* and *in vitro* following transfer into *Ae*. *aegypti*. An increase in cholesterol sequestration to lipid droplets occurs in *w*Mel / wMelPop-carrying *Ae*. *aegypti* cells, and treatment with the cyclodextrin 2HPCD released this stored cholesterol and induced a partial recovery of DENV replication (17). However, it is unclear if these changes occur for all virus-inhibiting strains of *Wolbachia*.

Release programs using *Wolbachia*-carrying *Ae*. *aegypti* for dengue transmission control are underway in a number of countries (2,18,19), using strain *w*Mel or strain *w*AlbB originating in *Aedes albopictus*. An intervention trial using *w*AlbB in Malaysia showed 40-80% reduction in dengue incidence over multiple release sites (18). With the continued field deployment of *Wolbachia* it is increasingly important to understand the molecular mechanisms underlying *Wolbachia*-mediated antiviral activity. Knowledge of the viral inhibition mechanisms will allow more informed monitoring and mitigation of potential operational problems, such as the possibility of viral resistance mutations that confer resistance to or the instability of particular strains of the symbiont in given environments. When *Ae*. *aegypti* larvae are reared at temperatures above ~35°C the density and maternal transmission of *w*Mel is lowered - potentially compromising its capacity to inhibit dengue in hot conditions and elevating the risk of selection of escape mutations (8,20–24). If *Wolbachia* strains can be identified for use in release programs that have mechanistic differences to *w*Mel / *w*AlbB in their viral inhibition, this would be highly valuable for long-term success of the strategy, in providing a means to either reduce the risk of selection of viral escape mutations, and / or allow a means of mitigation against viral escape should it occur.

In light of the unusually efficient viral inhibition conferred by strain *w*Au, which does not seem to be a consequence solely of its relatively high intracellular density (8,16), we sought to examine whether any differences could be identified relative to other *Wolbachia* strains in terms of the viruses inhibited or the cellular perturbations that may underlie virus inhibition. Cellular and proteomic analyses were utilized to compare the effects of *Wolbachia* strains in *Ae*. *aegypti* cell culture and dissected midgut tissues (17).

## Results

### Lipid perturbation and cyclodextrin profile

Lipid metabolism and transport are important for the inhibition of DENV in mosquito cells containing *w*Mel / *w*MelPop (17). In line with a previous study, cholesterol dynamics and effects of treatment with the cyclodextrin 2HPCD were carried out in *Ae*. *aegypti* midguts and *Ae*. *albopictus* cells containing either *w*Au, *w*Mel or negative for *Wolbachia*. In contrast to observations with *w*Mel, *w*Au-carrying midguts (Figure 1 a) and cells did not show an accumulation of cholesterol in lipid droplets (as seen by distinct punctate spots), and 2HPCD in cells led to a significant increase in lipid accumulation (Figure 1b & c). Zika replication was not rescued in *w*Au-carrying cells treated with 2HPCD, but was rescued in *w*Mel cells, except for the highest concentration (Figure 1d) - similar to previous observations for DENV although interestingly here we observed complete rescue of ZIKV replication (17).

**Figure 1.**
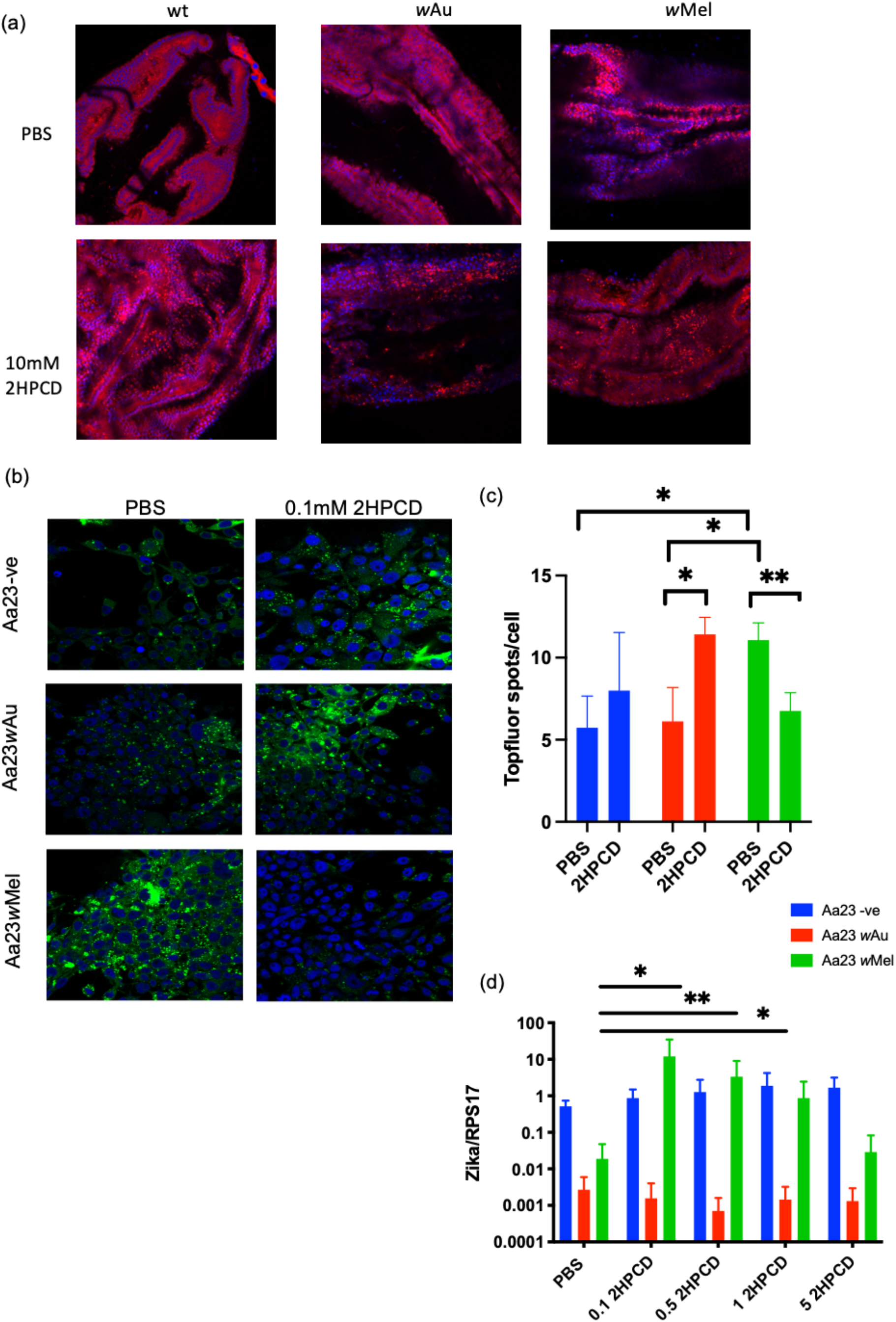
Effect of 2HPCD treatment on cholesterol dynamics in *Aedes aegypti* midguts and *Aedes albopictus* cell lines and on Zika replication. (a) *Aedes aegypti* mosquitoes were injected with 10 mM of 2HPCD or PBS, left to recover for 2 days before a bloodmeal was given. 72hr post bloodmeal midguts were dissected, fixed and stained with Nile red (red) and DAPi (blue) to detect intracellular lipid droplets (distinct punctate staining) and cell nuclei respectively. (b) Cell were pulse labelled for 30 mins with Topfluor (green), a fluorescent cholesterol derivative and treated for 48hr in either 2HPCD or PBS in order to measure cholesterol dynamics. Cell nuclei were stained with DAPi (blue). (c) Cellprofiler was used to calculate the number of Topfluor spots per cell in 3 independent replicates for each treatment (d) Cells were treated for 48hr with either PBS or 2HPCD and then infected with Zika at an MOI of 1. 72hr post infection total RNA was isolated. Data is normalised to the mosquito gene RPS17. Each bar corresponds to 3 independent experiments with 2 biological replicates per experiment (* P< 0.05 show significant differences in comparisons between the PBS control and each 2HPCD concentration for a given *Wolbachia* infection status, Mann-Whitney)

### *w*Au induces distinct changes in protein expression

The density of *Wolbachia* strains such as *w*Au and *w*Mel has been shown to correlate with the ability to block viruses (15,16). However, the strain *w*AlbA is found in similar densities and with similar tissue tropism to *w*Au, but shows relatively low anti-viral activity (8,16). Therefore, these strains can be compared to wildtype (wt) mosquitoes to determine mechanisms related to antiviral activity independent of density. In order to gain further insight into the mechanisms of *w*Au antiviral activity, a proteomic analysis was carried out on age-matched female midguts of *w*Au, *w*AlbA and wt *Ae*. *aegypti*. Midguts were chosen as previous proteomic analysis had shown results obtained from these tissues were robust for the study of *Wolbachia*/viral interactions (17). In total, 3821 proteins were detected, of which 27 were identified as *Wolbachia* proteins, which were subsequently excluded from the KEGG pathway analysis. From the total proteins identified, 3379 were quantified in all sample groups and were therefore used for differential expression analysis.

A principal component analysis performed on protein expression levels generated a clear separation of biological replicates according to *Wolbachia* status / strain (Figure 2a), differences in protein expression profiles could also be readily visualised in a heatmap representation of quantified proteins (Figure 2b). A linear modelling-based approach to differential expression analysis detected the greatest level of dysregulation in *w*Au/wt with 1088 significantly different proteins, followed by *w*Au/*w*AlbA with 765 dysregulated proteins and *w*AlbA/wt with 706 dysregulated proteins at 5% FDR (Figure 2c, Supplementary dataset). Volcano plot analysis of the differentially expressed proteins within each comparison shows a clear distinction between both *Wolbachia* strains and wt midguts and between the two strains, indicating clear proteome perturbations in the presence of *Wolbachia*. As expected, *Wolbachia* proteins are amongst the most significantly different in comparisons to uninfected midguts, 19 *Wolbachia* proteins could be detected as significantly elevated compared to the uninfected control. Relative abundance of the *Wolbachia* proteins detected above background indicate similar densities of *w*Au and *w*AlbA (Figure 2c), confirming the suitability of this system to study density independent differences in the level of antiviral activity between *Wolbachia* strains.

**Figure 2.**
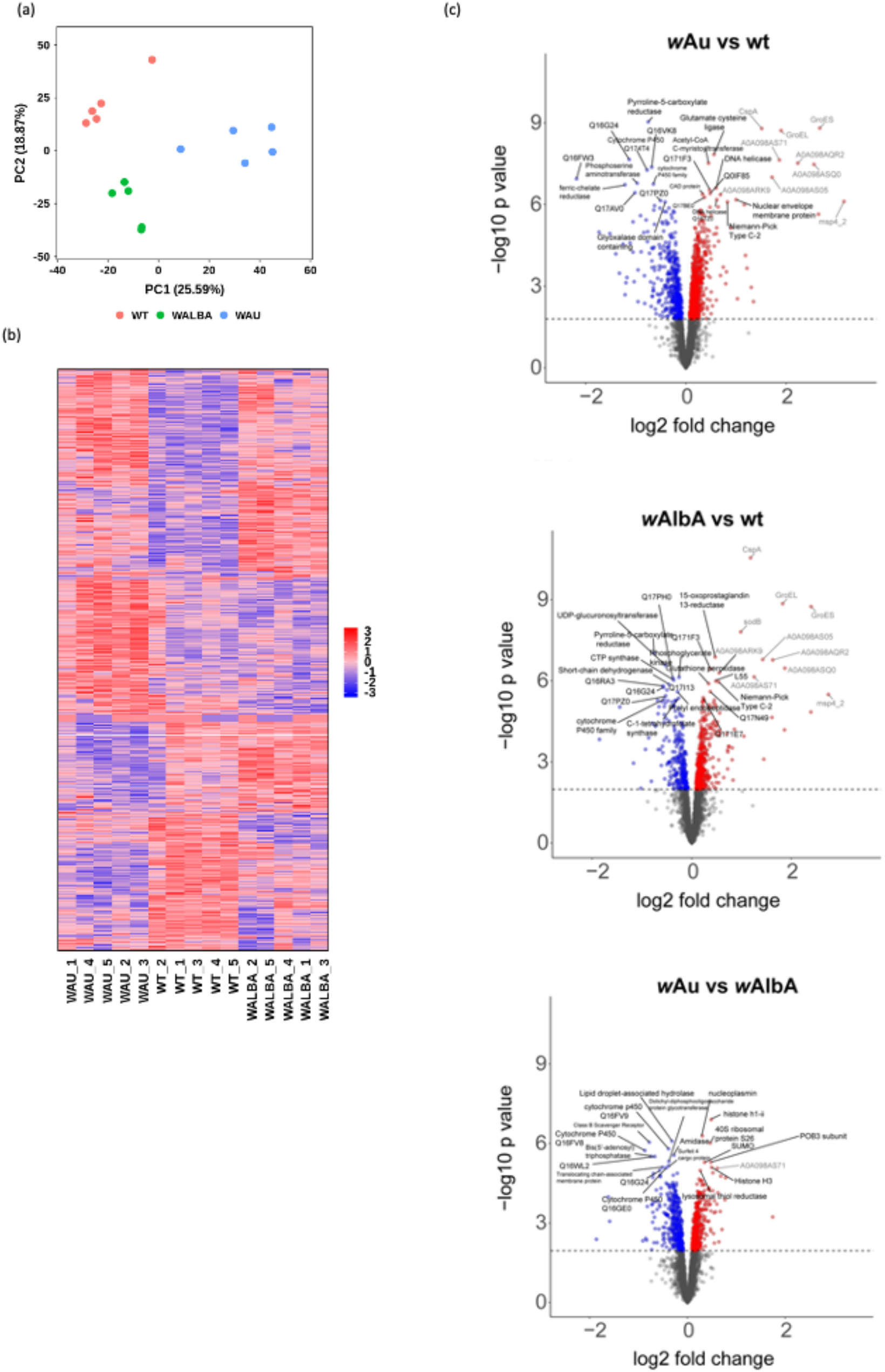
Comparison of proteins differentially expressed in *w*Au, *w*AlbA and wt midguts of *Aedes aegypti*. Pooled midguts were analysed by mass spectrometry to determine proteins that show significant alterations in expression between midguts containing *w*Au, *w*AlbA and wt (uninfected) N=5. (a) Principal component analysis of all quantified proteins, each dot represents a single replicate. (b) Clustered heatmap of significantly dysregulated proteins when *w*Au or *w*AlbA are present compared to wt. (c) mass spectrometry quantitation of 3379 proteins identified from midguts of *Aedes aegypti*. Proteins present at significantly different levels are highlighted in red (up regulated) or blue (down regulated), dashed line denotes p-value threshold for a false discovery rate of 5%. Top 20 differentially regulated *Aedes* proteins are labelled, with most enriched *Wolbachia* proteins in grey

### Comparison to *w*Mel proteomics data

As lipid perturbations appear different in *w*Au, the proteome dataset was compared to a previous dataset obtained for *w*Mel transinfected midguts, focusing on proteins hypothesised or known to affect antiviral activity in *w*Mel (17). The ER/unfolded protein response and lipid metabolism were previously shown to be markedly perturbed in midguts containing *w*Mel (17). Perturbations in proteins associated with lipid homeostasis were not seen in the *w*Au set (Table S1), where *w*Au is consistently different to *w*Mel. Upregulation of proteins involved in the protein unfolding response and increases in ER stress proteins were observed in the presence of *w*Mel; however, the opposite was observed for *w*Au (Table S1), with downregulation of some of the proteins that were upregulated in the *w*Mel line, and no significant difference in others. Interestingly, the *w*AlbA line showed changes broadly intermediate between *w*Mel and *w*Au. Taken together these data show that *w*Au differs from *w*Mel with regard to effects on lipid metabolism.

Since there is a large dynamic response to *Wolbachia*, a global analysis was undertaken using the StringDB database (25) in order to identify dysregulated pathways. The *Wolbachia*-carrying lines were compared to examine all proteins significantly dysregulated relative to the corresponding wildtype midguts. Proteins significantly dysregulated in *w*Au infected midguts relative to *w*AlbA infected midguts were also examined. A KEGG pathway analysis was conducted to examine the significantly over-represented pathways amongst the differentially regulated host proteins in each *Wolbachia* type, our previously published dataset comparing *w*Mel and uninfected midguts was analysed as previously (17)(Figure 3a). Several pathways known to be involved in *Wolbachia* growth and metabolism were dysregulated in all *Wolbachia*-carrying lines, as expected. Genes involved in fatty acid synthesis and amino acid synthesis are absent from the genome of *Wolbachia*; therefore these processes are likely dysregulated when the bacterium is present (26–30). The upregulation of proteasome proteins may be indicative of the need for a controlled breakdown of host proteins to increase amino acid availability (31). However, there were marked differences between the three *Wolbachia*-carrying lines and in particular, between *w*Au and *w*Mel. The *w*Au containing midguts have a broader profile of pathway dysregulation compared to the *w*AlbA and *w*Mel midguts, for example affecting lysosomes, DNA replication and glycan degradation. To separate out host cellular pathway alterations specifically associated with antiviral activity of *Wolbachia* from the generalised perturbations caused by the presence of the bacterium, *Ae*. *aegypti* proteins significantly dysregulated in *w*Au midguts relative to *w*AlbA midguts were analysed for enrichment of KEGG pathways (Figure 3a). This effectively subtracted the ‘background’ perturbations to show changes more likely to be specifically associated with *Wolbachia* antiviral activity, revealing a much more limited set of perturbed pathways. N-Glycan biosynthesis, protein processing in endoplasmic reticulum, ribosome biogenesis, protein export and endocytosis are significantly dysregulated in *w*Au compared to *w*AlbA midguts which are all known to be important for viral replication. Two pathways involving RNA, RNA transport and the spliceosome were significantly affected which may directly affect the ability of viral RNA to replicate efficiently in the cell.

**Figure 3.**
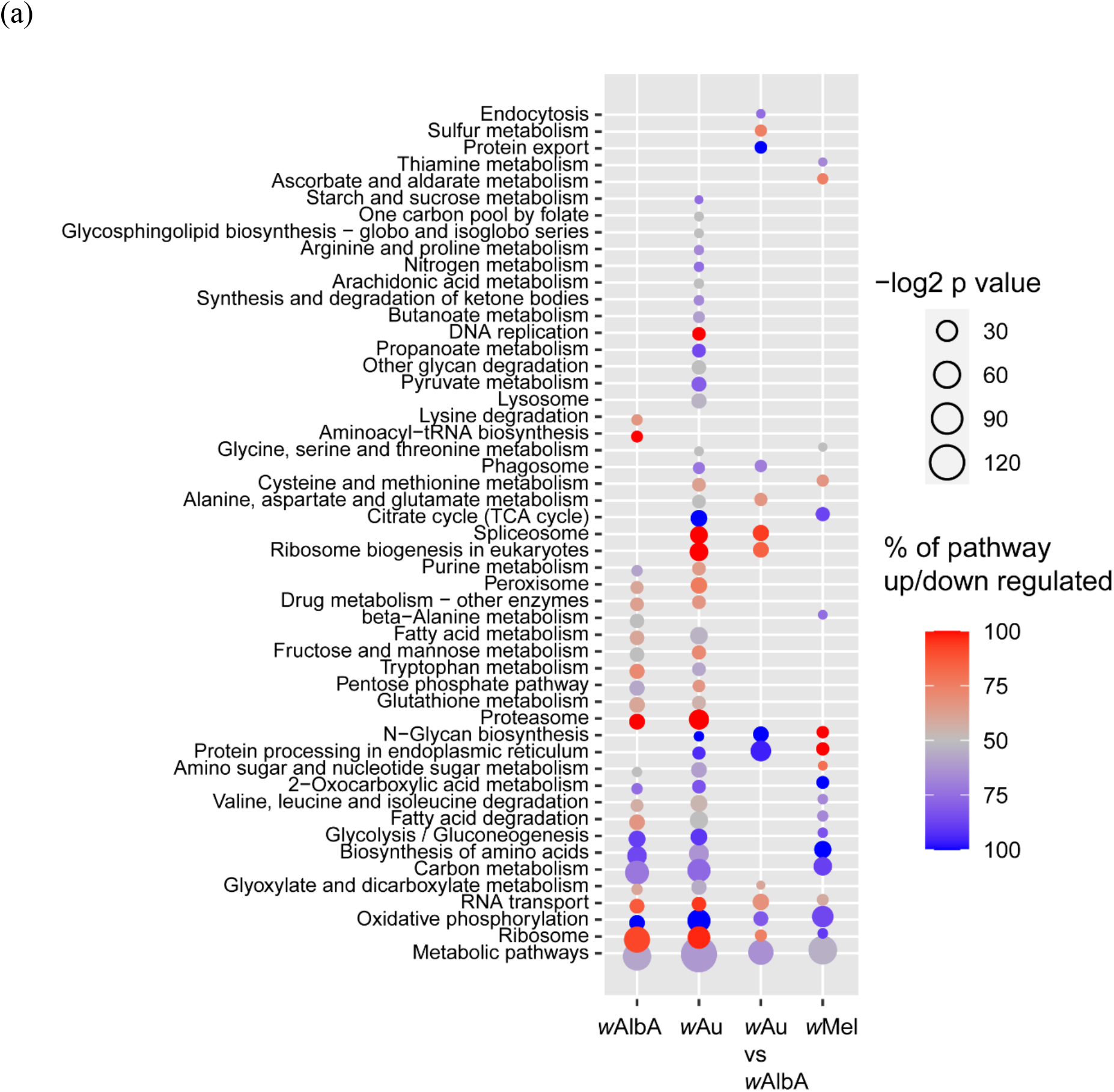
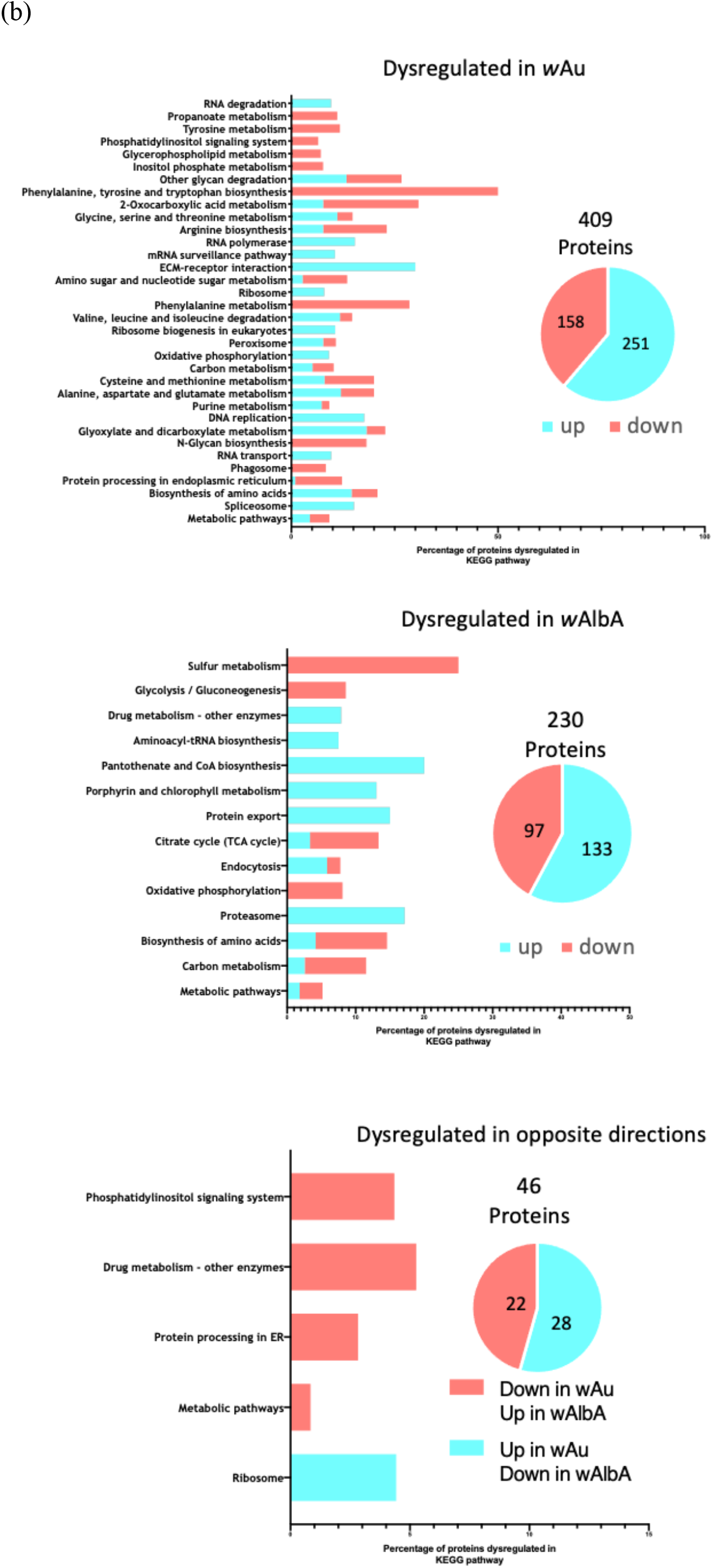
KEGG pathway analysis of differentially regulated proteins & pathway enrichment analysis based on *Wolbachia* strain-specific changes. (a) Significantly dysregulated host proteins (FDR <5%) were used to calculate over-represented KEGG pathway terms. Dysregulation is relative to uninfected midguts except the *w*Au vs *w*AlbA comparison. Bubble size is proportional to the statistical significance of the KEGG pathway enrichment. Proportion of each KEGG pathway up or down regulated is indicated with red or blue respectively. (b) Proteins significantly dysregulated between *w*Au and *w*AlbA when compared to wt midguts were split into 3 criteria and KEGG pathway analysis carried out in order to determine pathways significantly enriched in each criterion (FDR adjusted p-value <0.05).

### Dynamics of *w*Au and *w*AlbA dysregulation of proteins and subdivision into criteria

To further characterise pathways that may be responsible for the greater antiviral activity associated with *w*Au compared to *w*AlbA, additional KEGG pathway enrichment analyses were carried out, this time by subdividing into 3 criteria: proteins that are either specifically dysregulated with *w*Au, specifically dysregulated with *w*AlbA or dysregulated in both strains but in opposite directions. More proteins were found to be upregulated compared to downregulated in the first two criteria. A considerably higher number of proteins were found to be significantly dysregulated specifically in the *w*Au line (409 proteins and 35 KEGG pathways) compared to *w*AlbA (230 proteins and 14 KEGG pathways), suggesting that *w*Au has a greater impact on the host proteome (Figure 3b). Since the density of these two strains is similar within the midguts (4), this demonstrates that host perturbations are not solely determined by *Wolbachia* density. Few proteins were found to be dysregulated in opposite directions: 22 proteins were found to be downregulated in *w*Au but upregulated in *w*AlbA, while 28 were upregulated in *w*Au and downregulated in *w*AlbA.

Of key interest, given the differing levels of inhibition between the *w*Au and *w*AlbA lines, were KEGG pathways enriched only in the *w*Au line, and those found to be oppositely dysregulated when *w*Au and *w*AlbA are compared; perturbed pathways already known to be involved in DENV replication are discussed below.

### RNA pathways and translation initiation are specifically enriched in *w*Au

Since DENV and other RNA viruses rely host cell machinery to replicate, RNA pathway disruption may be important for *Wolbachia* mediated antiviral activity (32,33). Of the 35 KEGG pathways enriched only in *w*Au, 8 pathways are associated with RNA, DNA and splicing. Of the proteins dysregulated in these pathways all are upregulated in the *w*Au line compared to wildtype midguts. For comparison these proteins are not significantly dysregulated with *w*AlbA and show differing results in *w*Mel - for example, ribosomal proteins are downregulated with *w*Mel. These results indicate that there is a marked increase in RNA processing activity with *w*Au compared to the *Wolbachia*-negative line and lines carrying the *w*AlbA and *w*Mel strains.

Analysis of protein binding domain enrichment shows that only *w*Au has an enrichment of proteins with RNA binding domains such as ribonuclear protein La (AAEL003664) and the RNA binding protein Musashi (AAEL008257), again indicating an upregulation of pathways involved in RNA processing (Table S2). By comparing the pathways opposingly dysregulated in *w*Au and *w*AlbA midguts, it can be clearly seen that ribosome biogenesis is significantly enriched in *w*Au but significantly downregulated in *w*AlbA.

### ER functions & trafficking

As highlighted in Table S1 *w*Au and *w*AlbA lines show alterations in proteins associated with ER function and the unfolded protein response - important pathways in viral replication - that differ from *w*Mel. The *w*Au line shows reduced glucuronosyltransferase and glycosyltransferase activity compared to *w*AlbA. Further affected processes involving trafficking and protein processing included the ECM receptor pathway, which was significantly upregulated in the *w*Au line but not with the other two strains. In mosquitoes this pathway is known to be involved in the stability of the extracellular matrix in the midgut and may play a role in the midgut infection barrier (34). Of the pathways specific to *w*AlbA the only pathways that are not also enriched in *w*Mel are endocytosis and protein export, both of which are involved in arbovirus entry and replication/assembly.

## Discussion

It is important to determine how *Wolbachia* density affects levels of antiviral activity protection and indeed whether dysregulation of any given gene / protein could be involved in antiviral activity simply because it increases *Wolbachia* density. However it is possible to have a high density *Wolbachia* strain that shows comparatively low antiviral activity, and lipid metabolism manipulations can reverse antiviral activity without affecting *Wolbachia* density (8,17). It is currently unclear if the mechanisms of antiviral activity are conserved across *Wolbachia* strains and indeed the full extent of *Wolbachia*’s ability to block any RNA virus. The strain *w*Au has been shown to produce high levels of antiviral activity against several viruses; however, the mechanisms behind this are currently unknown. *w*Mel is known to have a profound effect on lipid and cholesterol metabolism and storage in both midguts and cell lines carrying the strain. In turn, manipulation of cholesterol storage by the cyclodextrin 2HPCD, leads to a partial rescue of DENV replication in cells (17).

Proteomic analysis of the *w*Au and *w*AlbA *Ae*. *aegypti* lines and the comparison to *w*Mel shows a clear disruption of metabolic pathways when compared to wt. For example, fatty acid synthesis and proteasome pathways are significantly dysregulated, supporting the hypothesis that *Wolbachia* manipulates these pathways to maintain itself within the host cell (26–30). The diet and metabolic state of insect cells is known to affect the density of *Wolbachia* and therefore there are likely key interactions between host metabolism and *Wolbachia* (27,35,36). The *w*Au and *w*Mel lines showed striking differences in pathways previously demonstrated to have a role in antiviral activity. Cholesterol and lipid metabolism alterations seen in the *w*Mel line at the proteomic level were not observed with *w*Au, and the accumulation of cholesterol in lipid droplets that occurs with *w*Mel was not seen in *w*Au-carrying midguts or cultured cells. The ability of 2HPCD to rescue *Wolbachia* antiviral activity in cells infected with Zika virus also differed markedly between strains - *w*Mel treated with 2HPCD for 48hrs resulted in increased Zika replication, but this was not the case for *w*Au-carrying cells.

The results presented here indicate that there are significant mechanistic differences underlying antiviral activity between *Wolbachia* strains. The presence of different mechanisms suggests that, should virus resistance evolve to counteract a particular strain of the symbiont, this resistance may not necessarily function against all *Wolbachia* strains. Therefore, the sequential use of multiple strains with different antiviral mechanisms could be a method to prevent or counteract viral evolution. The ability of arboviruses to adapt and escape *Wolbachia* antiviral activity has not been reported (37,38) but is a potential risk to the long-term efficacy of *Wolbachia* dengue control programs. Another possibility is that *Wolbachia* strains may adapt in the field to have lower density or restricted tropism in the host. Experiments using field-caught strains of *Ae*. *aegypti* carrying *w*Mel or *w*AlbB that have been under field selection for extended periods, found that both strains maintained high levels of transmission inhibition when challenged with dengue virus (2,39), but longer-term monitoring is needed.

KEGG pathway analyses revealed that a number of pathways involved in RNA biogenesis, translation and RNA recognition were significantly dysregulated in *w*Au cells. Given the requirement for DENV and other arboviruses to replicate using host machinery, the disruption of these pathways is noteworthy. Flaviviruses in particular are known to hijack the RNA degradation and surveillance pathways in order to replicate (40). The RNA binding protein Musashi, upregulated in *w*Au, for example is known to bind the 3’UTR of Zika virus and prompt replication/translation, and has been linked to pathogenicity (41). The ribonuclear protein La (AAEL003664), is upregulated in the *w*Au line, downregulated for *w*AlbA and not significantly dysregulated for *w*Mel. DENV infection causes a re-localisation of the protein and it is found to inhibit replication in a dose dependent manner (42). The RNA decay pathway has been implicated in antiviral activity of *w*Mel; it has been shown that *w*Mel*-*mediated antiviral activity against DENV in *Ae*. *aegypti* cells can be reduced by decreasing the levels of XRNI, a key protein involved in RNA decay (43).There is no increase in XRNI expression associated with *w*Mel, suggesting a functional change rather than a simple increase in protein availability and thus degradation of viral RNA. Therefore, there is a suggestion that RNA decay may play a part in *w*Mel antiviral activity and assessing the effect of deletion of XRNI on *Wolbachia* density would be useful to further investigate this. XRNI depletion in *w*Au lines may also be of interest to determine the effect on antiviral activity, *Wolbachia* titres and the RNA decay pathway.

ER trafficking pathways, glycosyltransferases and protein processing in the ER, disrupted specifically by wAu, are required for DENV translation and folding of viral proteins. Dengue and other arboviruses do not encode glycosyltransferases, which are crucial to several aspects of the viral life cycle. Downregulation of these proteins in insect hosts can have a profound effect on virus binding, replication, protein folding and egress (44). Protein processing in the ER is likely to be significantly reduced in *w*Au with 12% of the pathway downregulated. DENV replication and assembly relies on these processes in the ER (45). In order to reach the ER for replication, DENV undergoes clathrin mediated endocytosis (46); interestingly AAEL014375, a clathrin coat assembly protein, is significantly downregulated in *w*Au but not in *w*Mel or *w*AlbA. While *w*Mel appears to affect DENV replication/translation due in part to a lack of available cholesterol / lipids, this does not seem to be the case for *w*Au. Comparison of virus localization after entry and the dynamics of viral replication may help clarify the mechanistic differences further. In *w*AlbA protein export and endocytosis were increased and this may facilitate virus entry. Very few pathways were found to be significantly enriched only in *w*AlbA compared to *w*Au.

Host transcriptome analysis in *Drosophila melanogaster* naturally carrying *w*Mel shows that nucleotide metabolism, RNA binding and processing and translation, and transcription are perturbed, similar to *w*Au in this study (47). This indicates that the host background is also important when determining host interactions and that assessing single tissues and whole organisms may produce different results. Interestingly *w*Mel also appears to have an effect on gene splicing (47) ; therefore as *w*Au perturbs the spliceosome it would be interesting to look at transcript profiles in *w*Au and *w*Mel-carrying mosquitoes. The clear evidence for differences between strains of *Wolbachia* in proteomic changes that are likely to affect virus inhibition presented here suggest that the virus inhibition of *w*Au could be caused by disruption in pathways involved in RNA biogenesis/processing. Differing mechanisms of *Wolbachia*-mediated antiviral activity means that it should be possible to create a panel of lines carrying different strains for field releases that could be deployed to restore efficacy in the event of viral evolution and escape mutations. The study provides a foundation for further functional investigations into the mechanisms by which different strains inhibit virus transmission.

## Materials and Methods

### Mosquito rearing and cell work

Mosquito colonies were maintained at standard 27°C and 70% relative humidity with a 12-hour light/dark cycle. Lines have been described previously (8,13). *Wolbachia*-free lines consist of the original line which *Wolbachia* was transferred into, giving all lines the same genetic background. Tetracycline cured lines were not used as the removal of other bacteria from the lines may have resulted in a skewed proteome not related to the presence or absence of *Wolbachia*. Further to this there is currently no data on the long-term effect of *Wolbachia* on midgut proteomes and if these changes persist after removal of *Wolbachia*. Age matched (4 days old), mosquitoes were injected in the thorax with 414nl of 10mM 2HPCD (this concentration was chosen in line with previously published *in vivo* experiments in mice and humans (48,49) using Nanoject II (Drummond Scientific, Pennsylvania, USA) hand-held microinjector, with a pulled glass capillary. 48hr after injection the mosquitoes were blood fed using a Hemotek artificial blood-feeding system (Hemotek, UK) using defibrinated sheep blood (TCS Biosciences, UK). Mosquitoes were allowed to recover for 72hr before midguts were dissected and stained as described below.

In Aa23 (*Aedes albopictus*) cells which had been cleared of *Wolbachia*, *w*Mel and *w*Au strains were introduced from *Drosophila simulans* STCP lines (50) as follows: Aa23 cells were plated the day before in a 96-well plate. For each *Wolbachia* strain to be transferred, around 200 mated *Drosophila* flies were placed in a BugDorm rearing cage (W17.5 x D17.5 x H17.5 cm) with a Petri dish containing grape agar (3% agar, 1% sucrose, 25% grape juice, water) and a spot of yeast paste in the centre to stimulate egg-laying. After one hour, around 500 *Drosophila* eggs were collected from the agar plate with a brush and rinsed in sterile water. Eggs were further dechorionated and surface-sterilized in 2.5% bleach for 2 min, 70% ethanol for 5 min twice and were rinsed in sterile water three times. Sterilized eggs were transferred to a 1.5 mL Eppendorf tube, resuspended in PBS and homogenized with a sterile pestle. The egg homogenate was centrifuged at 2,500 g for 10 min at 4°C to remove cellular debris and the supernatant was filtered through a 5 μm and a 2.7 μm Millex syringe filters. The filtered homogenate was finally centrifuged at 18,500 g for 5 min at 4°C to pellet the bacteria. The bacterial pellet was resuspended in 100 μl Schneider’s media with 10% FBS and overlaid onto the aa23 cells. Finally, the cell plate was centrifuged at 2,500g for 1h at 15°C. In the following days, fully confluent cells were serially passaged from the 96-well plate, to 48, 24 and 12-well plates. Cells were later maintained in 25 cm^3^ flasks with Schneider’s media with 10% FBS at 28°C. Cells were checked regularly for *Wolbachia* density using quantitative PCR as described previously (8).

### Virus infection in cells

For Zika infection, cells were plated out in 24 well plates at a density of 5×10^4^/ml and left to settle for 24hr. After 24hr various concentrations of 2HPCD or PBS was added at varying concentrations and incubated for 48hr for 48hr in Schneider’s media supplemented with 10% FCS. Media was then removed and Zika (PRVABC59-strain) added at a multicity of infection of 1 in fresh media. Media was then removed and Zika (PRVABC59-strain) added at a multicity of infection of 1. Cells were incubated for 72hr. Cells and media were collected at 24hr and 72hr post infection. Following the removal of media Trizol (ThermoFisher UK) was added, and RNA was extracted following manufactures protocol. cDNA was synthesised using the All-In-One cDNA Synthesis SuperMix (Biotools, TX, USA). ZIKV was quantified using ZIKV 835 and ZIKV 911c primers (ZIKV-835: TTGGTCATGATACTGCTGATTGC, ZIKV-911c: CCTTCCACAAAGTCCCTATTGC). Values were normalised to the RpS17 mosquito gene (Rps17-F: CACTCCCAGGTCCGTGGTAT, Rps17-R: GGACACTTCCGGCACGTAGT) as reference by relative expression (Pfaffl method (51)). qPCR was carried out on a Rotor Gene Q machine (Qiagen) using 2x qQuantiNova SYBR. The following program was used to run the qPCRs: 95 °C for 5 min, 40× 393 cycles of 95 °C for 15 s and 60 °C for 30 s, followed by a melt-curve analysis.

### Staining and imaging

Following dissection, midguts were fixed with Fixative solution (ThermoFisher UK (United Kingdom)) for 10 min, followed by 3 washes in PBS. Midguts were then incubated in Nile red (Sigma) stain at 0.1ug/ml for 40 min, followed by three PBS washes and mounted in ProLong™ Gold Antifade Mountant with DAPi (ThermoFisher UK). Aa23 cells were pulse labelled with the cholesterol derivative Topfluor as previously described (17) and treated as above with either 2HPCD (2 hydroxypropyl β cyclodextrin) or PBS. All images were then acquired using a Zeiss LSM 880 confocal microscope (Zeiss) with a 63X objective for cells and 20X objective for midguts. Nile red was imaged using a 488 nm laser. Nuclei stained with DAPI were imaged using a 405 nm laser detector. TopFluor was imaged using a 488 nm laser, with GaAsP detectors. All settings were obtained by first imaging uninfected Aa23 *Wolbachia*-negative cells incubated in PBS as a standard control. For midguts settings were obtained by first imaging wt, PBS as a standard control. Quantification was carried out by imaging 3 independent ×64 images from 3 independent wells on a 24-well optical plate. Images were analysed using Cell Profiler. A global image threshold was set using the Otsu method and images were analysed in order to identify the number of nuclei and the number of green spots corresponding to TopFluor staining. Data are presented as the number of spots per cell. Large crystals in cells due to precipitated Topfluor were masked from images to ensure only intracellular fluorescence was measured.

### Proteomics sample preparation

Proteomic analysis was carried out on midguts from age matched (10 days old), non-bloodfed, female mosquitoes. Each biological replicate consisted of pooled midguts from 20 individuals, 5 biological replicates were analysed for each *Wolbachia* infection type. Each biological replicate was lysed in 200μl 8M urea 50mM triethylammonium bicarbonate (TEAB) supplemented with 1x protease inhibitor (Roche). Midguts were sonicated for 3 cycles of 15 seconds yielding approximately 100μg of protein from each pool as measured by BCA assay (Thermo Scientific). Samples were reduced with 5mM DTT for 30 mins at 50°C then alkylated with 15mM IAA for 30 mins at RT. Urea was diluted to a final concentration of 1.5M and trypsin/Lys-C (Promega) added to a ratio of 25:1 protein:trypsin. After overnight digestion at 37°C, the digest was acidified with 0.5% trifluoroacetic acid (v:v) and centrifuged at 18,000g for 7 mins. Digested peptides were desalted using 50mg C18 cartridges (Phenomenex Strata) and dried down. Peptides were resuspended in 50mM TEAB and labelled with a TMT 10plex kit (Thermo Scientific). From each biological replicate, 6μg of peptide was taken, this was pooled and labelled with the 131 TMT channel to create a common pool reference channel enabling relative quantification across all 15 samples. 44μg of peptide from each biological replicate was labelled, replicates were pooled into two groups. Each group was fractionated by high pH reversed phase fractionation according to manufacturers instructions (Thermo Scientific).

### Mass spectrometry

Peptides from midgut samples were resuspended in 0.1% formic acid and loaded onto an UltiMate 3000 RSLCnano HPLC system (Thermo) equipped with a PepMap 100 Å C18, 5 μm trap column (300 μm × 5mm, Thermo) and a PepMap, 2 μm, 100 Å, C18 Easy Nanonanocapillary column (75 μm × 150 mm, Thermo). The trap wash solvent was 0.05% (v:v) aqueous TFA and the trapping flow rate was 15 μl/min. The trap was washed for 3 min before switching flow to the capillary column. Separation used gradient elution of two solvents: solvent A, aqueous 1% (v:v) formic acid; solvent B, aqueous 80% (v:v) acetonitrile containing 1% (v:v) formic acid. The flow rate for the capillary column was 300 nl/min and the column temperature was 40 °C. The linear multi-step gradient profile was: 3–10% B over 8 min, 10–35% B over 125 min, 35–65% B over 50 min, 65–99% B over 7 min and then proceeded to wash with 99% solvent B for 4 min. The column was returned to initial conditions and re-equilibrated for 15 min before subsequent injections.

The nanoLC system was interfaced with an Orbitrap Fusion hybrid mass spectrometer (Thermo) with an EasyNano ionisation source (Thermo). Positive electrospray ionisation (ESI)-MS, MS2 and MS3 spectra were acquired using Xcalibur software (version 4.0, Thermo). Instrument source settings were: ion spray voltage, 1,900 V; sweep gas, 0 Arb; ion transfer tube temperature, 275 °C. MS1 spectra were acquired in the Orbitrap with: 120,000 resolution, scan range: m/z 380–1,500; automatic gain control (AGC) target, 2e5; max fill time, 50 ms. Data-dependant acquisition was performed in top speed mode using a 4 s cycle, selecting the most intense precursors with charge states >1. Dynamic exclusion was performed for 50 s post-precursor selection and a minimum threshold for fragmentation was set at 3e4. MS2 spectra were acquired in the linear ion trap with: scan rate, turbo; quadrupole isolation, 1.2 m/z; activation type, collision-induced dissociation; activation energy: 35%; AGC target, 1e4; first mass, 120 m/z; max fill time, 50 ms. MS3 spectra were acquired in multi notch synchronous precursor mode (SPS3), selecting the 5 most intense MS2 fragment ions between 400 and 1,000 m/z. SPS3 spectra were measured in the Orbitrap mass analyser using: 50,000 resolution, quadrupole isolation, 2 m/z; activation type, HCD; collision energy, 65%; scan range: m/z 110–500; AGC target, 5e4; max fill time, 86 ms. Acquisitions were arranged by Xcalibur to inject ions for all available parallelisable time.

### MS data analysis

TMT data peak lists were converted from centroided .raw to .mgf format using Mascot Distiller (version 2.6.1, Matrix Science) and MS3 spectra were concatenated into their parent MS2 spectra for database searching. Mascot Daemon (version 2.5.1, Matrix Science) was used to combine .mgf files and search against a subset of the UniProt database containing Ae. aegypti and Wolbachia w Mel proteins (17,811 sequences) using a locally running copy of the Mascot program (Matrix Science Ltd, version 2.5.1). Search criteria specified: Enzyme, trypsin; Fixed modifications, Carbamidomethyl (C), TMT10plex (N-term, K); Variable modifications, Oxidation (M); Peptide tolerance, 5 p.p.m.; MS/MS tolerance, 0.5 Da; Instrument, ESI-TRAP. The Mascot .dat result file was imported into Scaffold Q + (version 4.7.5, Proteome Software) and a second search run against the same database using X!Tandem was run. Protein identifications were filtered to require a maximum protein and peptide FDR of 1% with a minimum of two unique peptide identifications per protein. Protein probabilities were assigned by the Protein Prophet algorithm. Proteins that contained similar peptides and could not be differentiated based on MS/MS analysis alone were grouped to satisfy the principles of parsimony. Proteins sharing significant peptide evidence were grouped into clusters. Quantification of relative protein abundance was calculated from TMT reporter ion intensities with Scaffold Q + using the common pool reference channel. TMT isotope correction factors were taken from the document supplied with the reagents by the manufacturer.

Normalised log2 transformed protein intensities were analysed with Limma (52)to determine significant differences between sample groups at a 5% False Discovery Rate, options ‘trend’ and ‘robust’ were enabled in the empirical Bayes procedure. Multiple testing correction was carried out according to Benjamini & Hochberg. For the KEGG pathway bubble plot, significantly dysregulated proteins detected by Limma were submitted to StringDB to detect over-represented pathways. Significantly regulated proteins from *w*AlbA (p<0.0135), *w*Au (p<0.01559), *w*Mel (p<0.01463) midguts were split into downregulated and upregulated groups for each *Wolbachia* type, each group was then submitted to StringDB to calculate enriched KEGG pathways (25). Further to this, for comparison of *w*Au and *w*AlbA, data was split into 3 criteria as outlined in Figure 5 for KEGG pathway analysis.

## Acknowledgments

We thank Adam Dowle and Tony Larson in the Technology Facility at the University of York for carrying out mass spectrometry analysis of samples and providing expertise.

## References

1. Fraser JE, De Bruyne JT, Iturbe-Ormaetxe I, Stepnell J, Burns RL, Flores HA, et al. Novel *Wolbachia*-transinfected *Aedes aegypti* mosquitoes possess diverse fitness and vector competence phenotypes. PLoS Pathog. 2017 Dec;13(12):e1006751.

2. Hoffmann AA, Montgomery BL, Popovici J, Iturbe-Ormaetxe I, Johnson PH, Muzzi F, et al. Successful establishment of *Wolbachia* in *Aedes* populations to suppress dengue transmission. Nature. 2011 Aug 24;476(7361):454–7.

3. Joubert DA, Walker T, Carrington LB, De Bruyne JT, Kien DHT, Hoang NLT, et al. Establishment of a *Wolbachia* Superinfection in *Aedes aegypti* Mosquitoes as a Potential Approach for Future Resistance Management. PLoS Pathog. 2016 Feb;12(2):e1005434.

4. Bhatt S, Gething PW, Brady OJ, Messina JP, Farlow AW, Moyes CL, et al. The global distribution and burden of dengue. Nature. 2013 Apr 25;496(7446):504–7.

5. Puntasecca CJ, King CH, LaBeaud AD. Measuring the global burden of chikungunya and Zika viruses: A systematic review. PLOS Neglected Tropical Diseases. 2021 Mar 4;15(3):e0009055.

6. Moreira LA, Iturbe-Ormaetxe I, Jeffery JA, Lu G, Pyke AT, Hedges LM, et al. A *Wolbachia* symbiont in *Aedes aegypti* limits infection with dengue, Chikungunya, and Plasmodium. Cell. 2009 Dec 24;139(7):1268–78.

7. Walker T, Johnson PH, Moreira LA, Iturbe-Ormaetxe I, Frentiu FD, McMeniman CJ, et al. The *w*Mel *Wolbachia* strain blocks dengue and invades caged *Aedes aegypti* populations. Nature. 2011 Aug 24;476(7361):450–3.

8. Ant TH, Herd CS, Geoghegan V, Hoffmann AA, Sinkins SP. The *Wolbachia* strain *w*Au provides highly efficient virus transmission blocking in *Aedes aegypti*. PLoS Pathog. 2018;14(1):e1006815.

9. Bian G, Xu Y, Lu P, Xie Y, Xi Z. The Endosymbiotic Bacterium *Wolbachia* Induces Resistance to Dengue Virus in *Aedes aegypti*. PLOS Pathogens. 2010 Apr 1;6(4):e1000833.

10. Rainey SM, Martinez J, McFarlane M, Juneja P, Sarkies P, Lulla A, et al. *Wolbachia* Blocks Viral Genome Replication Early in Infection without a Transcriptional Response by the Endosymbiont or Host Small RNA Pathways. PLOS Pathogens. 2016 Apr 18;12(4):e1005536.

11. Schultz MJ, Tan AL, Gray CN, Isern S, Michael SF, Frydman HM, et al. *Wolbachia w*Stri Blocks Zika Virus Growth at Two Independent Stages of Viral Replication. mBio. 2018 May 22;9(3).

12. Blagrove MSC, Arias-Goeta C, Di Genua C, Failloux A-B, Sinkins SP. A *Wolbachia w*Mel transinfection in *Aedes albopictus* is not detrimental to host fitness and inhibits Chikungunya virus. PLoS Negl Trop Dis. 2013;7(3):e2152.

13. Blagrove MSC, Arias-Goeta C, Failloux A-B, Sinkins SP. *Wolbachia* strain *w*Mel induces cytoplasmic incompatibility and blocks dengue transmission in *Aedes albopictus*. PNAS. 2012 Jan 3;109(1):255–60.

14. Lu P, Bian G, Pan X, Xi Z. *Wolbachia* induces density-dependent inhibition to dengue virus in mosquito cells. PLoS Negl Trop Dis. 2012;6(7):e1754.

15. Martinez J, Tolosana I, Ok S, Smith S, Snoeck K, Day JP, et al. Symbiont strain is the main determinant of variation in *Wolbachia*-mediated protection against viruses across Drosophila species. Molecular Ecology. 2017;26(15):4072–84.

16. Chouin-Carneiro T, Ant TH, Herd C, Louis F, Failloux AB, Sinkins SP. *Wolbachia* strain *w*AlbA blocks Zika virus transmission in Aedes aegypti. Med Vet Entomol. 2020;34(1):116–9.

17. Geoghegan V, Stainton K, Rainey SM, Ant TH, Dowle AA, Larson T, et al. Perturbed cholesterol and vesicular trafficking associated with dengue blocking in *Wolbachia*-infected *Aedes aegypti* cells. Nature Communications. 2017 Sep 13;8(1):526.

18. Nazni WA, Hoffmann AA, NoorAfizah A, Cheong YL, Mancini MV, Golding N, et al. Establishment of *Wolbachia* Strain *w*AlbB in Malaysian Populations of Aedes aegypti for Dengue Control. Current Biology. 2019 Dec 16;29(24):4241–4248.e5.

19. Tantowijoyo W, Andari B, Arguni E, Budiwati N, Nurhayati I, Fitriana I, et al. Stable establishment of *w*Mel *Wolbachia* in *Aedes aegypti* populations in Yogyakarta, Indonesia. PLOS Neglected Tropical Diseases. 2020 Apr 17;14(4):e0008157.

20. Hoffmann AA, Iturbe-Ormaetxe I, Callahan AG, Phillips BL, Billington K, Axford JK, et al. Stability of the *w*Mel *Wolbachia* Infection following Invasion into *Aedes aegypti* Populations. PLoS Negl Trop Dis [Internet]. 2014 Sep 11 [cited 2020 Nov 19];8(9). Available from: https://www.ncbi.nlm.nih.gov/pmc/articles/PMC4161343/

21. Mancini MV, Ant TH, Herd CS, Gingell DD, Murdochy SM, Mararo E, et al. High temperature cycles result in maternal transmission and dengue infection differences between *Wolbachia* strains in *Aedes aegypti*. bioRxiv. 2020 Nov 26;2020.11.25.397604.

22. Ross PA, Ritchie SA, Axford JK, Hoffmann AA. Loss of cytoplasmic incompatibility in *Wolbachia*-infected *Aedes aegypti* under field conditions. PLoS Negl Trop Dis [Internet]. 2019 Apr 19 [cited 2020 Nov 19];13(4). Available from: https://www.ncbi.nlm.nih.gov/pmc/articles/PMC6493766/

23. Ross PA, Wiwatanaratanabutr I, Axford JK, White VL, Endersby-Harshman NM, Hoffmann AA. *Wolbachia* Infections in *Aedes aegypti* Differ Markedly in Their Response to Cyclical Heat Stress. PLOS Pathogens. 2017 Jan 5;13(1):e1006006.

24. Ulrich JN, Beier JC, Devine GJ, Hugo LE. Heat Sensitivity of *w*Mel *Wolbachia* during *Aedes aegypti* Development. PLOS Neglected Tropical Diseases. 2016 Jul 26;10(7):e0004873.

25. Szklarczyk D, Gable AL, Lyon D, Junge A, Wyder S, Huerta-Cepas J, et al. STRING v11: protein-protein association networks with increased coverage, supporting functional discovery in genome-wide experimental datasets. Nucleic Acids Res. 2019 08;47(D1):D607–13.

26. Jiménez NE, Gerdtzen ZP, Olivera-Nappa Á, Salgado JC, Conca C. A systems biology approach for studying *Wolbachia* metabolism reveals points of interaction with its host in the context of arboviral infection. PLOS Neglected Tropical Diseases. 2019 Aug 30;13(8):e0007678.

27. Molloy JC, Sommer U, Viant MR, Sinkins SP. *Wolbachia* Modulates Lipid Metabolism in *Aedes albopictus* Mosquito Cells. Appl Environ Microbiol. 2016 May 15;82(10):3109–20.

28. Pietri JE, DeBruhl H, Sullivan W. The rich somatic life of *Wolbachia*. MicrobiologyOpen. 2016 Dec;5(6):923–36.

29. Voronin D, Bachu S, Shlossman M, Unnasch TR, Ghedin E, Lustigman S. Glucose and Glycogen Metabolism in *Brugia malayi* Is Associated with *Wolbachia* Symbiont Fitness. PLOS ONE. 2016 Apr 14;11(4):e0153812.

30. Wu M, Sun LV, Vamathevan J, Riegler M, Deboy R, Brownlie JC, et al. Phylogenomics of the reproductive parasite *Wolbachia pipientis w*Mel: a streamlined genome overrun by mobile genetic elements. PLoS Biol. 2004 Mar;2(3):E69.

31. Fallon AM, Witthuhn BA. Proteasome activity in a naïve mosquito cell line infected with *Wolbachia pipientis w*AlbB. In Vitro Cell Dev Biol Anim. 2009 Sep;45(8):460–6.

32. Paranjape SM, Harris E. Control of dengue virus translation and replication. Curr Top Microbiol Immunol. 2010;338:15–34.

33. Walsh D, Mohr I. Viral subversion of the host protein synthesis machinery. Nat Rev Microbiol. 2011 Oct 17;9(12):860–75.

34. Dong S, Behura SK, Franz AWE. The midgut transcriptome of Aedes aegypti fed with saline or protein meals containing chikungunya virus reveals genes potentially involved in viral midgut escape. BMC Genomics [Internet]. 2017 May 15 [cited 2020 Nov 20];18. Available from: https://www.ncbi.nlm.nih.gov/pmc/articles/PMC5433025/

35. Kremer N, Voronin D, Charif D, Mavingui P, Mollereau B, Vavre F. *Wolbachia* Interferes with Ferritin Expression and Iron Metabolism in Insects. PLoS Pathog [Internet]. 2009 Oct 23 [cited 2020 Nov 20];5(10). Available from: https://www.ncbi.nlm.nih.gov/pmc/articles/PMC2759286/

36. Voronin D, Cook DAN, Steven A, Taylor MJ. Autophagy regulates *Wolbachia* populations across diverse symbiotic associations. PNAS. 2012 Jun 19;109(25):E1638–46.

37. Koh C, Audsley MD, Di Giallonardo F, Kerton EJ, Young PR, Holmes EC, et al. Sustained *Wolbachia*-mediated blocking of dengue virus isolates following serial passage in *Aedes aegypti* cell culture. Virus Evol [Internet]. 2019 Jan 1 [cited 2020 Nov 20];5(1). Available from: https://academic.oup.com/ve/article/5/1/vez012/5512689

38. Martinez J, Bruner-Montero G, Arunkumar R, Smith SCL, Day JP, Longdon B, et al. Virus evolution in *Wolbachia-infected Drosophila*. Proc Biol Sci [Internet]. 2019 Nov 6 [cited 2020 Nov 20];286(1914). Available from: https://www.ncbi.nlm.nih.gov/pmc/articles/PMC6823055/

39. Noor Afizah A, Mancini M-V, Ant TH, Martinez J, Kamarul G, Nazni W, et al. *Wolbachia* strain *w*AlbB maintains high density and dengue inhibition following introduction into a field population of *Aedes aegypti*. Philosophical Transactions of the Royal Society B: Biological Sciences [Internet]. 2020 Aug 5 [cited 2020 Nov 20]; Available from: http://eprints.gla.ac.uk/221898/

40. Akiyama BM, Eiler D, Kieft JS. Structured RNAs that evade or confound exonucleases: function follows form. Curr Opin Struct Biol. 2016 Feb;36:40–7.

41. Chavali PL, Stojic L, Meredith LW, Joseph N, Nahorski MS, Sanford TJ, et al. Neurodevelopmental protein Musashi-1 interacts with the Zika genome and promotes viral replication. Science. 2017 Jul 7;357(6346):83–8.

42. García-Montalvo BM, Medina F, del Angel RM. La protein binds to NS5 and NS3 and to the 5’ and 3’ ends of Dengue 4 virus RNA. Virus Res. 2004 Jun 15;102(2):141–50.

43. Thomas S, Verma J, Woolfit M, O’Neill SL. *Wolbachia*-mediated virus blocking in mosquito cells is dependent on XRN1-mediated viral RNA degradation and influenced by viral replication rate. Vignuzzi M, editor. PLoS Pathog. 2018 Mar 1;14(3):e1006879.

44. Idris F, Muharram SH, Diah S. Glycosylation of dengue virus glycoproteins and their interactions with carbohydrate receptors: possible targets for antiviral therapy. Arch Virol. 2016 Jul;161(7):1751–60.

45. Reid DW, Campos RK, Child JR, Zheng T, Chan KWK, Bradrick SS, et al. Dengue Virus Selectively Annexes Endoplasmic Reticulum-Associated Translation Machinery as a Strategy for Co-opting Host Cell Protein Synthesis. Journal of Virology [Internet]. 2018 Apr 1 [cited 2020 Oct 26];92(7). Available from: https://jvi.asm.org/content/92/7/e01766-17

46. Cruz-Oliveira C, Freire JM, Conceição TM, Higa LM, Castanho MARB, Da Poian AT. Receptors and routes of dengue virus entry into the host cells. FEMS Microbiol Rev. 2015 Mar 1;39(2):155–70.

47. Lindsey AR, Bhattacharya T, Hardy RW, Newton IL. *Wolbachia* and virus alter the host transcriptome at the interface of nucleotide metabolism pathways [Internet]. Microbiology; 2020 Jun [cited 2021 Jan 13]. Available from: http://biorxiv.org/lookup/doi/10.1101/2020.06.18.160317

48. Matsuo M, Togawa M, Hirabaru K, Mochinaga S, Narita A, Adachi M, et al. Effects of cyclodextrin in two patients with Niemann-Pick Type C disease. Mol Genet Metab. 2013 Jan;108(1):76–81.

49. Tanaka Y, Yamada Y, Ishitsuka Y, Matsuo M, Shiraishi K, Wada K, et al. Efficacy of 2-Hydroxypropyl-β-cyclodextrin in Niemann–Pick Disease Type C Model Mice and Its Pharmacokinetic Analysis in a Patient with the Disease. Biological and Pharmaceutical Bulletin. 2015;38(6):844–51.

50. Martinez J, Longdon B, Bauer S, Chan Y-S, Miller WJ, Bourtzis K, et al. Symbionts commonly provide broad spectrum resistance to viruses in insects: a comparative analysis of *Wolbachia* strains. PLoS Pathog. 2014 Sep;10(9):e1004369.

51. Pfaffl MW. A new mathematical model for relative quantification in real-time RT–PCR. Nucleic Acids Res. 2001 May 1;29(9):e45.

52. Ritchie ME, Phipson B, Wu D, Hu Y, Law CW, Shi W, et al. limma powers differential expression analyses for RNA-sequencing and microarray studies. Nucleic Acids Research. 2015 Apr 20;43(7):e47–e47.

